# On eco-evolutionary dynamics of phenologies in competitive communities and their robustness to climate change

**DOI:** 10.1101/2021.12.01.470711

**Authors:** Thomas Cortier, Nicolas Loeuille

**Affiliations:** iEES, Institut of ecology and environmental sciences of Paris, Pierre and Marie Curie campus, 4 place Jussieu 75005 Paris

**Keywords:** Phenology, community ecology, evolutionary rescue, competition, adaptive dynamics, dispersal

## Abstract

**G**lobal changes currently cause temporal shifts in the favourable conditions for different phases of species life cycles. Phenologies characterizing temporal presence, may adapt through heritable evolution in response to these changes. Given a community context, this evolution may cause a change in the phenology overlap and thus a change of interspecific interactions such as competition. Using a model in which phenologies compete and coevolve, we study the conditions under which diversity emerges, as well as their annual distribution. We find that the environment richness (food quantity, light, pollinators…) and competition constrain the diversity and spread of phenologies. A robust pattern of phenologies distribution emerges consistent with Swedish flowering observations. Once a stable community is reached, we apply a progressive change in environmental conditions. We found that adaptation eventually restored diversity, but that the simulated change often led to numerous extinctions due to increased competition. The percentage of diversity lost depends on the speed of change and on the initial diversity. Phenologies already pre-adapted to the new environmental conditions drive the restoration of diversity after the change. We finally study a spatial version of the model in which local communities are organized along an environmental gradient. Pre-change, allowing dispersal decreases the local adaptation of phenologies to their local fixed environmental conditions. Dispersal however largely enhances the maintenance of biodiversity in changing environments, though its benefits are not homogeneous in space. Evolution remains the only rescue mechanism for southern phenotypes.

## Introduction

Brittany, once a humid region favoring artichoke cultivation may well soon become a new French wine-growing region. On the other hand, in traditional wine-growing regions, grapes ripen earlier in the season. Such agricultural shifts immediately pinpoint how species phenologies are currently affected by climate changes. Wild species undergo similar changes and many recent shifts in phenology and distributions have been documented (Parmesan and Yohe 2003).

Phenologies are simultaneously selected by environmental conditions and interspecific interactions. Depending on the species life cycle and phenotypic traits, some parts of the year may be more favorable for investment in growth, reproduction and resource acquisition. Environmental factors favoring the presence of a species can be linked to fluctuations in temperature, light or to the availability of certain limiting nutrients (J. Lloyd and Taylor 1994). For a given species, the combination of favorable abiotic conditions can be seen as a temporal niche space or environmental window. The presence of other species however constrains species phenology on an ecological and evolutionary scale. Within a functional group, species present at the same time may find themselves in competition for resources or space. Flowering plants may be in competition for pollinators (Mitchell et al. 2009), while developing amphibian larvae may compete with breeding species within a pond (Alford and Wilbur 1985). Mutualistic and trophic interactions similarly require overlapping phenologies for the two species (Loeuille 2019; Waser 1979), so that the fitness of partner species largely depends on their relative timing. Interspecific relationships thereby likely influence the coevolution of species phenologies in natural communities.

Competition can cause phenologies to diversify. Assuming a suitable temporal window, phenotypes active at different times partly avoid competition. Niche partitioning and limiting similarity is then expected, so that a given diversity of phenologies may coexist (Macarthur and Levins 1967; Chesson 2000). Such competitive effects are for instance well known for flowering species. In the Arima Valley of Trinidad (tropical region), fruiting seasons of the 19 species of Miconia genus are spread out to cover the entire year. The staggered fruiting season would have evolved through interspecific competition (Snow 1965). An annual temporal partitioning of pollinators has also been reported many times as an important selective factor of flowering phenologies (Appanah 1985; Botes, Johnson, and Cowling 2008; Wheelwright 1985).

Under current global changes, environmental conditions largely vary in space (warmer conditions northward and at higher altitude) and in time (eg, earlier onset of phenologies, (Sherry et al. 2007; Vitousek 1992)). Such changes in the abiotic conditions directly affect the environmental window in which phenologies may coevolve. Evolution of phenological traits or dispersal to colonize new parts of the changing environmental niche can both save a population. Consistent with this view, many instances of rapid evolution of phenologies have been recently reported (Carter, Saenz, and Rudolf 2018; Franks, Sim, and Weis 2007; Husby, Visser, and Kruuk 2011; Jonzen 2006; Nussey 2005; Phillimore et al. 2010). Earlier arrival dates of migratory birds have for instance been noted, with evolution likely playing a key role (Jonzen 2006). If evolution is fast enough to keep a match between the species phenology, its interactor and the environmental window, evolutionary rescue is expected (Gomulkiewicz and Holt 1995). As global changes also shift the fundamental niche of species in space, dispersal of phenologies to match the environmental niche could similarly offer a rescue mechanism. Northward shifts and phenological shifts have been simultaneously noted in many groups of species (Guisan and Thuiller 2005; Parmesan and Yohe 2003). The extent of dispersal or local adaptation depends on the species ability to disperse and on the environmental fragmentation (Kubisch et al. 2013; Loeuille 2019). For warm latitudes (equator, tropics), no source of phenotypes adapted to warmer temperature may exist. When the environment changes, local adaptation is then the only mechanism capable of saving biodiversity (Norberg et al. 2012; Sinervo et al. 2010).

Changes in the environment are not simply abiotic and climatic, but also entail variations in the community and interaction context. Reshuffling of interactions and local extinctions largely restructure present-day communities (Tylianakis et al. 2008). As matching of phenologies underpin ecological interactions, the restructuring of natural communities largely impacts the fitness of coexisting phenologies and can cause diversity losses. For instance, (Carter, Saenz, and Rudolf 2018) studied the phenology of 66 amphibian species on a 15 year interval. They showed that phenology overlap increased and proposed that this may increase competitive interactions by up to 25%. Eco-evolutionary consequences are intriguing. From an ecological point of view, the increased competition may lead to species loss, due to competitive exclusion. From an evolutionary point of view, given an increase in competitive selective pressures, further displacement of phenologies is expected. Note that the two aspects are in fact intertwined. If species have different abilities to evolve (population size, genetic diversity…), evolutionary rescue may save species that evolve, but increase competition on other species of the community (Loeuille 2019). Such asymmetric effects could make certain species more vulnerable and cause their extinction (Johansson 2008). Interspecific competition can also accentuate the selection gradient in certain situations and accelerate the evolution (Osmond and de Mazancourt 2013). Finally, when local diversity is large, niche packing constrains the possible evolution in response to environmental changes (de Mazancourt, Johnson, and Barraclough 2008). Changing competitive interactions in a community can therefore alter evolutionary rescue by changing the ability of species to track environmental changes.

We build a model of a community of competing phenologies in a temporal niche potential which we will call an environmental window. The environmental window is first fixed, then altered to mimic global changes. Phenologies are characterized by two traits, the spread and the mean date. Fitnesses of the different phenologies depend not only on abiotic constraints (environmental window) but also on surrounding phenologies (competition). Using this model, we investigate the following questions. (1) Given a fixed environmental window, and starting with just one phenology, we investigate the conditions of diversification. When it occurs, we characterize both the ecological (total diversity, community structure) and the evolutionary state (phenology distribution). Phenological traits distribution is then compared to an empirical dataset of flowering plants from Sweden. (2) We then study the impact of a change of the environmental window on traits and on local diversity. We link the loss of phenological trait diversity with the ability of species to evolve for both date and spread. We expect greater losses of biodiversity when evolutionary potential are decreased. (3) Finally, we undertake metacommunity simulations along a continuum of latitudes and investigate whether dispersal or phenology evolution better explain evolutionary rescue events.

## I Construction of the eco-evolutionary model

### I.1 Population dynamics of a given phenology (*μ_i_*, *σ_i_*)

For a given phenotype (*μ_i_*, *σ_i_*), population density *n_i_* follows a simple Lotka-Volterra system. Assuming that *k* phenologies coexist in the environment:

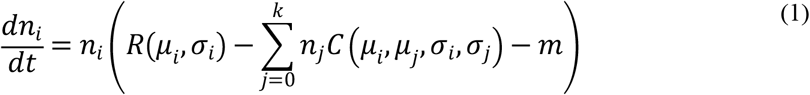

Here, *R* embodies the *per capita* reproduction rate and *C* the competition kernel, explained below. All parameters are detailed in table 1.

**Table 1 :** Model variables and their associated units.

| Variable | Significance | Unit |
| --- | --- | --- |
| $n_i$ | Population density of the $i^{\text{th}}$ phenotype phenology | $n$ |
| $\mu_i$ | Average date of the $i^{\text{th}}$ phenotype phenology | $date$ |
| $\sigma_i$ | Spread of the $i^{\text{th}}$ phenotype<br>phenology in days | *days* |
| $R(\mu_i, \sigma_i)$ | Intrinsic reproduction rate of the<br>population | $t^{-1}$ |
| $C(\mu_i, \mu_j, \sigma_i, \sigma_j)$ | Per capita competition rate<br>between phenotypes i and j | $t^{-1}.n^{-1}$ |

**Table 2 :** The parameters of the model, their meaning and their associated unit. The values reported are those used in the simulations. The intervals correspond to the variation of this parameter for the analyses.

| Parameter | Significance | Unit | Value |
| --- | --- | --- | --- |
| $m$ | The threshold for intrinsic reproduction to reach a<br>overall positive reproduction rate | $t^{-1}$ | 200000 |
| G | Ability of a species to exploit the environment | $days.t^{-1}$ | 200 |
| $P_{max}$ | Richness of the environment | $t^{-1}$ | [3;64] |
| C | Competition | $t^{-1}.n^{-1}$ | 0.01 |
| $\lambda$ | The slope of the resource function at the inflection<br>point. | $date^{-1}$ | 0.1 |
| $j_{1/2}$ | The date of the inflection of the sigmoid. | *date* | [60;120] |
| $P_{dep}$ | Richness depletion in summer | $t^{-1}$ | [0;0.9$P_{max}$] |
| $\lambda_{dep}$ | The slope of inflection | $date^{-1}$ | 0.5 |
| $j_{1/2dep}$ | The date of inflection of the depletion. | *date* | [170;120] |
| $\beta_\mu, \beta_\sigma$ | Poisson expectation of mutations per individual every<br>$dt_{mut}$ for $\mu$ and $\sigma$ | $n_{mut}.n^{-1}$ | [0;10]*$10^{-6}$ |
| $dt_{mut}$ | The time steps between mutations | $t$ | 0.1 |
| $T_{ch}$ | The duration of environmental change | $t$ | [40;9000] |
| $\omega_\mu, \omega_\sigma$ | The variance of the mutation distribution for $\mu$ and $\sigma$ | *date, days* | 0.1,0.01 |
| $r_{disp}$ | Poisson expectation of dispersers per individual every<br>$dt_{disp}$ | $n_{disp}.n^{-1}$ | [$10^{-7}$; $10^{-4}$] |
| $dt_{disp}$ | The time step between dispersal events | $t$ | 0.01 |

### I.2 Phenological traits and associated trade-offs

Phenological traits (*μ_i_*, *σ_i_*) define a temporal occupancy function *g*, modeled using a gaussian function centered on *μ_i_* (Equation 2, Figure 1a). Note that this function is normalized, so that when activity is spread on a larger time window (higher *σ*), it is also reduced (fig 1c):

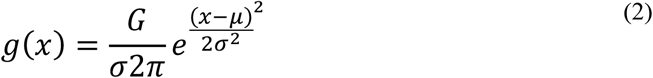

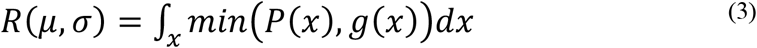

**Figure 1 :**
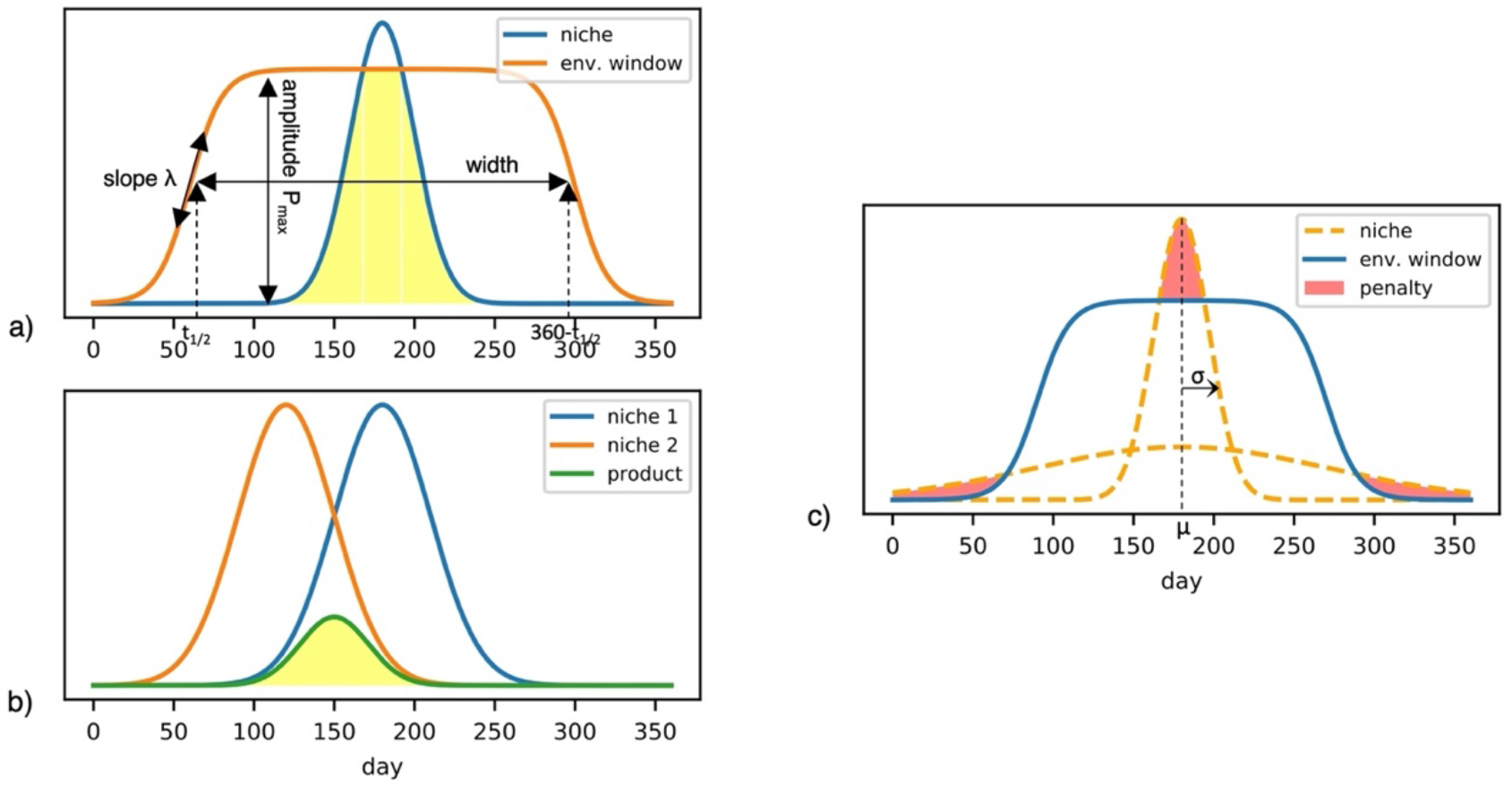
Defining the population dynamics of phenologies : a) The temporal occupancy function (equation 2, blue) and environmental window (equation 4, orange) define the intrinsic reproduction rate (equation 3, yellow integral). b) Competition between two phenologies (yellow integral) increases with their similarity (equation 5). c) Note that because of our definition of the reproduction rate, phenologies that are too narrow or too wide (variations in *σ*) pay a cost here represented by the red integral.

The match between the temporal occupancy function and the environmental window defines the *per capita* reproduction rate R (equation 3, fig1a). As displayed on figure 1c, this creates a trade-off acting on the spread *σ*, linking generalism and environmental use. When phenologies are spread on little time (small *σ*), the quality of the environment may not be sufficient to support this intense activity (narrow gaussian function overshoots the environmental window). However, plants may swamp their herbivores by producing vulnerable organs in concentrated bursts (Augspurger 1981). For large *σ* (spread phenology) on the other hand, energy may be too limiting at extreme dates. Exploiting an environment to reproduce can be resource-intensive (Primack and Stacy 1998).

Note that the environmental window (equation 4, fig 1a) summarizes biotic and abiotic conditions (excluding competition) suitable for the development of the considered phenotype. Such factors include temperature, rainfall, and resource availability. Taking the example of plants, vegetative phenologies often depend on the photoperiod, humidity and temperature (Chuine and Régnière 2017). Flowering phenology is constrained by the presence of pollinators and resource accumulation (Rathcke and Lacey 1985) while fruiting phenology may depend on the temporal availability of seed dispersers (Wheelwright 1985). We consider several scenarios for the environmental window. First, the environmental niche is fixed and made of two stuck sigmoids (first part of equation 4, fig 1A) and we let emerge the community from the diversification of phenologies. Suitability is maximum in the middle of the year at 180.

In equation 4, *P_max_* represents the quality of the environment. Parameter *j*_1/2_ is inversely related to the width of the environmental window. Parameter *λ* affects the steepness of the environmental window early and late in the year.

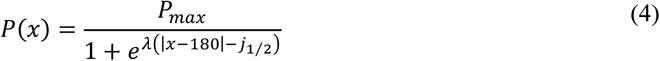

### I.3 Linking phenologies and competitive interactions

When phenologies co-occur within the environmental window, they are assumed to compete for available resources. Following classical frameworks (MacArthur and Levins 1964), we assume that competition is larger when phenologies are more similar, so that competition is proportional to the integral of the product of the two presence functions (equation 5).

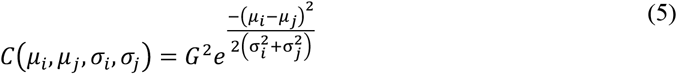

Note that individuals having exactly the same phenology compete at a maximum and constant rate *G^2^*.

## II Ecological dynamics

Consider first that all individuals have the same phenology. Equation (1) then simply corresponds to a logistic equation and the population reaches its carrying capacity (6).

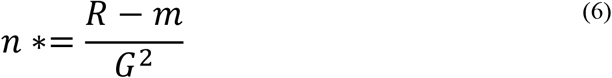

The population then exists whenever the intrinsic reproduction rate is higher than intrinsic mortality rate *m*.

## III Evolution, coevolution and diversification of phenologies in a fixed environment

Evolution of phenologies is studied using a numerical implementation of adaptive dynamics (Dieckmann and Law 1996). Mutations occur independently, either on the phenological spread *σ* or on the mean date *μ*.

The equations of population dynamics (equation 1) are integrated, and at fixed time intervals *dt_mut_*, the number of mutants is drawn in a poisson distribution that depend on the *per capita* mutation probability (β_μ_ for *μ, β_σ_* for *σ*) and on the size of the populations currently present in the community. While we typically consider rare mutations, we allow variations in the frequency of mutation *dt_mut_* to modulate the evolutionary potential of evolving phenologies. Whenever a population mutates, a rare mutant population is introduced (at density 1). Mutations are of small amplitude and the mutant trait is drawn in a normal distribution centered on the parent trait. The variance of this normal distribution is the parameter for *μ* and *ω_σ_* for *σ*. When integrating equation (1) any population passing below the extinction threshold (<1) is assumed extinct and dropped from the system.

While we are able to give a few mathematical results for simpler cases (see below), most scenarios are not mathematically trackable so that we mostly rely on simulations. A benefit of this simulation choice is that it allows us a large flexibility as we can make the environmental window take very diverse forms to investigate climate change, but also systematically vary the relative speed of ecology and evolution.

To uncover how the evolution of each trait (*μ*, *σ*) depends on the value of the other trait, but also on other parameters of the system, we first fix the environment and study the evolution of each trait separately, the other being fixed.

### III.1 Evolution and diversification of *μ*

We start with a single resident population of phenology (μ,σ) and consider the evolution of the mean date *μ*. The relative fitness of a mutant *μ*′ that differs slightly from the resident population *μ* is given by its invasion fitness (equation 7), ie, its growth rate when rare and the resident at equilibrium (Metz, Nisbet, and Geritz 1992):

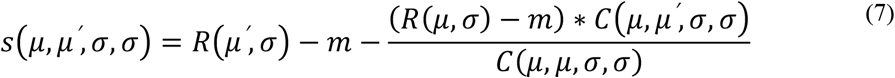

Because we assume no depletion, the first term of the equation will be maximum mid-year (creating stabilizing selection). On the contrary, the last term will be larger when phenological niches have less overlap, i.e. when *μ*′ differs from *μ*. This will favor disruptive selection, creating diversification when the environmental window is wide enough or when the spread *σ* is small enough.

Since the maximum of the environmental window is mid-year (x=180), the penalty of a wide or narrow phenological niches (fig 1c) will be minimal at this date. Evolutionary dynamics of the trait can be visualized using pairwise invasibility plots (PIPs, (Geritz et al. 1998)). A PIP (figure 2b) shows the sign of the invasive fitness (equation 7) of a mutant *μ*′ (y-axis) in a resident population *μ* (x-axis). Black areas show positive values where corresponding mutants invade the resident population. The direction of evolution can then be deduced (blue trajectories, red trajectories correspond to failed invasions). On figure 2b (left), note that for narrow phenologies (*σ* = 30), evolution eventually leads to a singularity at which both earlier and later mean dates are able to invade (blue dots at the end of the trajectory). Such situations correspond to branching points that allow diversification around *μ* =180 (fig2a). Because the environmental window is much larger than the phenology here, the fitness landscape is rather flat (large black areas along the diagonal from 100 to 260). Conversely, for wide phenologies (fig 2b, right), evolution leads to a unique singular strategy at *μ* =180 that cannot be invaded (read crosses). Such situations are called CSS (Continuously Stable Strategies, (Eshel 1983)) and are indicative of stabilizing selection. Evolution stops there and no diversification occurs.

**Figure 2 :**
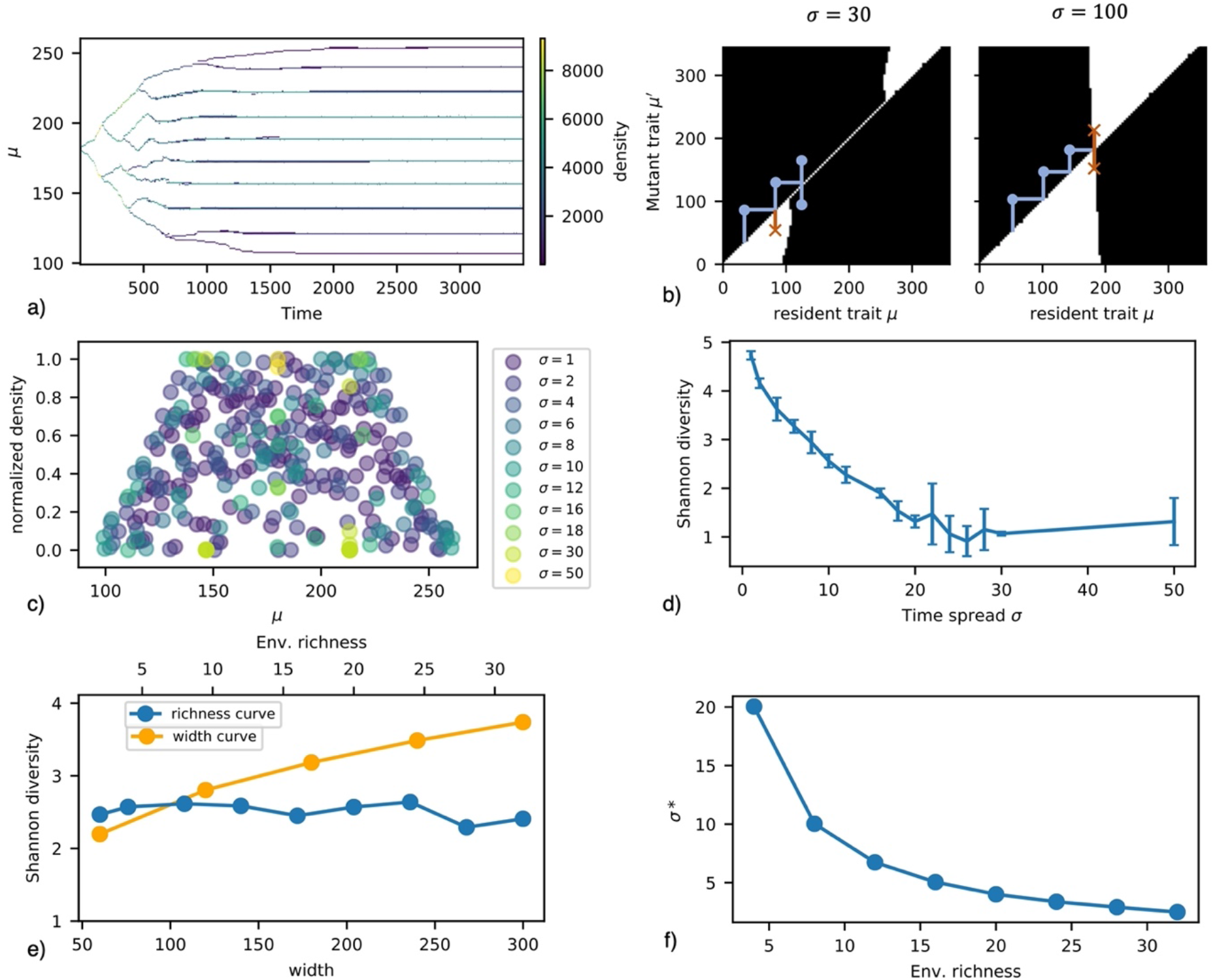
Evolution of the phenology mean date (panels a-e) and spread (panel f) depending on the environmental conditions and the other trait. a) Diversification of *μ* in time (*σ* = 10) b) Pairwise invasibility plot (PIP) showing the evolutionary dynamics of the mean date for two (fixed) values of *σ* (black: mutant *μ*′ can invade resident *μ*, white: mutant *μ*′ cannot invade resident *μ*). Example evolutionary trajectories are in blue while examples of failed invasions are in red. (*P_max_* = 3) c) Distribution of equilibrium phenology mean dates for several *σ* values. d) Shannon diversity at equilibrium according to *σ* with *μ* evolving (the error bars are the standard deviation for 10 replicates). e) Diversity of phenologies at equilibrium depending on the characteristics of the environmental window (amplitude and width) with *μ* evolving. f) Stable evolutionary equilibrium for phenology spread (*σ*) depending on the richness of the environmental window. (β_*μ*_, β_σ_ = 10^−5^ for panels (acdef), *j*_1/2_ = 120 for panels (acdf))

We now detail further situations in which diversification occurs. Figure 2c shows the quasi-equilibrium situations for different sets of simulations where spread is systematically manipulated. Not that the wider phenology (*σ* = 50) has only one date (yellow) and makes the limit beyond which no diversification occurs. When the spread is more limited (*σ* = 30, light green), the system eventually settles at two phenological dates (close to 150 and 210). Smaller phenological spreads lead to increasingly more phenological niches with heterogeneous densities. We quantified diversity using Shannon indices to account for density asymmetries. The equilibrium diversity of phenologies increases when the (fixed) phenological spread is smaller (figure 2d), which is consistent with niche packing expectations (Macarthur and Levins 1967).

Note that diversity also depends on characteristics of the environmental window (fig2e). While it varies rather little with the maximum richness of the environmental window *P_max_* (blue), it increases with the width of the environmental window (orange). Intuitively, given the phenological spread *σ* is fixed, it is possible to pack more phenologies when a wider environmental window exists.

### III.2 *σ* is evolving to accommodate the richness of the environment

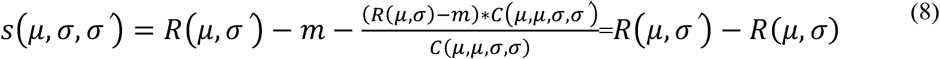

We now fix the phenology mean date *μ* and study the evolution of phenological spread *σ*. As before, evolution can be understood_by analyzing the invasion fitness of a rare mutant *σ*′ in a resident population *σ*:

Remembering that mutants can invade when this quantity is positive, equation 8 means that evolution will simply follow variations of the intrinsic reproduction rate. Penalties shown on figure 1c then simply have to be minimized. To minimize the first penalty, for a given date, the optimal phenology should match the environmental quality, not overshoot the window. Such a perfect match however happens for a given *σ* that may induce penalties early and late in the year. This last penalty can often be neglected if the phenology is not too spread or too close from the edge of the window. The optimal strategy under these conditions is stable and can be explicitly computed:

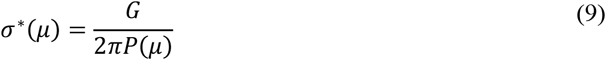

Equation 9 shows that the selected spread is inversely proportional to the quality of the environment, given the (fixed) mean phenological date *μ*. Phenologies are therefore more spread when the environment is poor (low *P_max_*). A direct consequence is that phenologies at the edge of the environmental window are expected to be more spread out. Numerical simulations confirm the mathematical analysis (figure 2f). We find that evolutionary equilibrium *σ* * decreases in richer environments, so that phenologies specialize in time.

We stress that no diversification is possible at *σ* *(CSS) so that in our model the diversification only happens for mean dates *μ*. This diversification is however constrained by both the environment and the spread of phenology *σ* (fig2d, e). The spread therefore potentially allows an adaptation to the abundance of the resource when *μ* has evolved. To confirm this, we now investigate the coevolution of the two traits.

### III.4 Coevolution of phenological traits

We now allow mutations to occur on both traits (*σ* and *μ*). We investigate how the shape of the environmental window affects the evolutionary equilibrium. An example simulation showing the emergence of seven different phenologies is displayed on fig3a. In link with previous analyses (equation 9), we observe that phenologies that evolve at central (high quality) dates evolve narrow phenologies, while phenologies evolving at extreme date become spread. This pattern of phenology distribution is robust to variations in the environmental window. For instance, on fig 3b, for nine different maximum qualities of the environment, we observe that the further the phenologies evolve from central dates, the greater their temporal spread (figure 3b). In summary, phenologies adapt to the environment richness at the date of their phenology which constrains the diversification indirectly through interspecific competition and the availability of ressources on both sides.

**Figure 3 :**
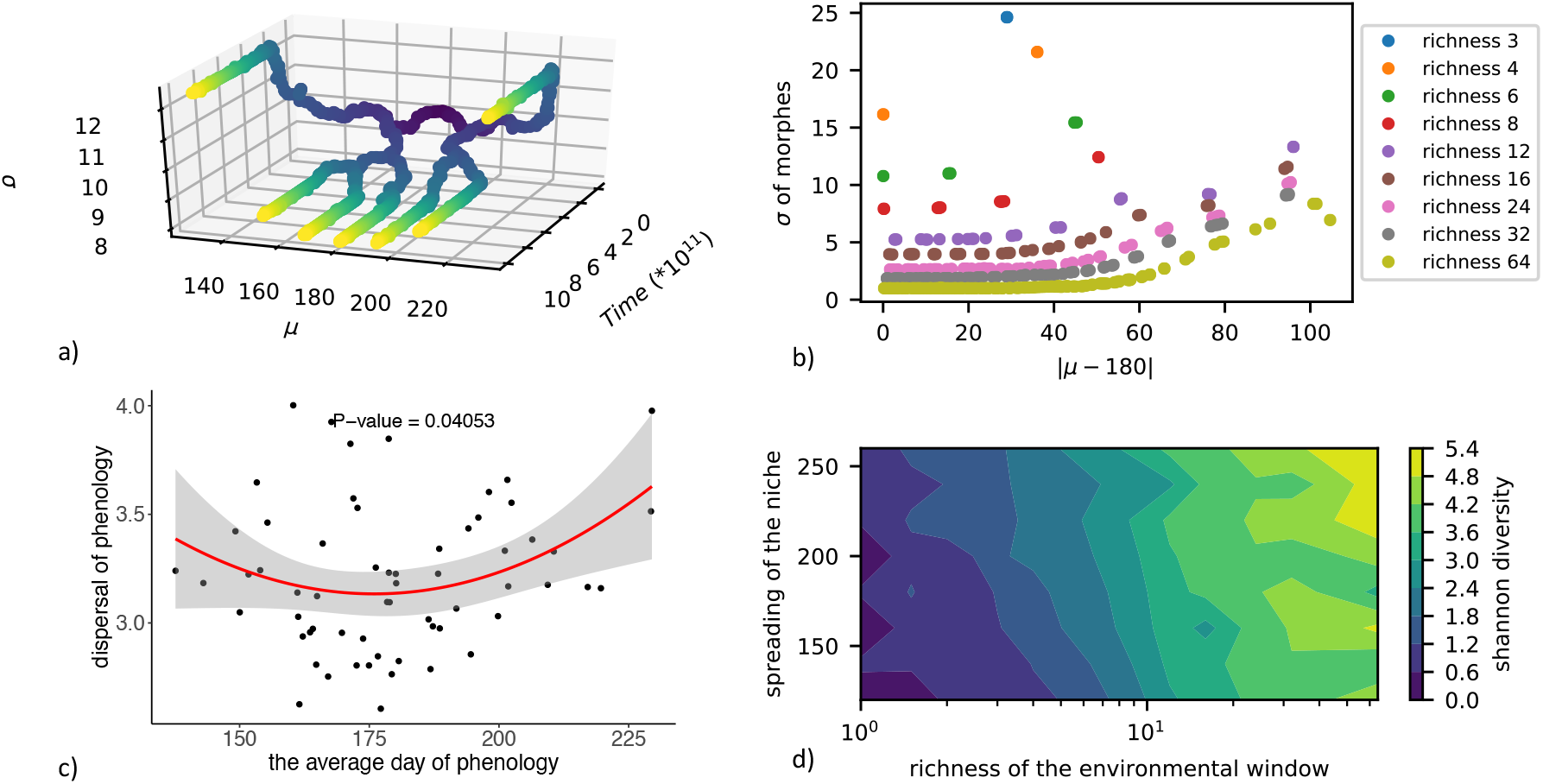
Coevolution in a stable environment and comparison with empirical data. a) Coevolutionary dynamics of the spread and average date of phenologies. The color yellows in time. (*P_max_* = 8) b) Distribution of evolved phenological spread *σ* depending on evolved mean date *μ* at equilibrium. Nine simulations are shown, for different qualities of the environmental window. In all cases, phenologies are more spread as the evolved dates become less central. c) Relationship between phenological spread and mean date for the flowering of 63 plant species in Sweden. d) Variations in Shannon diversity at coevolutionary equilibrium depending on the environmental window parameters. (β_*μ*_, β_σ_ = 10^−5^ for panels (abd), *j*_1/2_ = 120 for panels (ab))

Emerging diversity depends on the shape of the environmental window (figure 3d). Particularly, higher quality of the environmental window leads to higher diversity levels. The richer the environment, the more phenologies evolve to restricted temporal presence thereby avoiding competition with surrounding phenologies. The width of the environmental window here plays a minor role for diversity.

#### Empirical analysis of flowering phenological spread

We test whether the pattern of more spread phenologies at extreme dates (figure 3b) is relevant for species constrained by the same environmental window. We investigate how the flowering interval of 63 plant species in Sweden depends on their flowering time. For this dataset, the seasonality of the conditions favorable to the development of flowers (light, pollinators…) is very marked and reduced in time. 30 specimens per species were collected in the herbarium of the Swedish Natural History Museum (Weinbach 2015).

The number of flowers per specimen was reported for each observation. We fit a Gaussian on the counting of flowering observations through the year of each species. It is assumed that the sampling effort was the same for all species over time. Each sampling corresponds to a count of the number of flowers. Having count data, we use a generalized linear model with a poisson-like error:

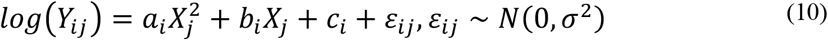

(10) *X_j_* being the day of the year, *Y_ij_* being the count of the flowering-species i on day j. *a_i_*, *b_j_* and *C_j_* are estimated for each species.

The mean date of *i* correspond to: 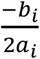

The temporal spread: 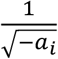

For the different species, we want to link the estimated spread to the estimated mean date. Equation (11) shows the statistical model with *X_i_* the mean date of species i and *Y_i_* its spread.

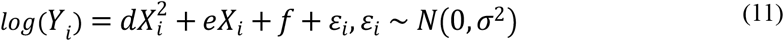

Consistent with our expectation, Figure 3c shows that plants flowering in the middle of the summer have shorter phenologies than those flowering at more extreme dates. Short spread flowering is more common in the summer when the competition might be stronger. An explanation could be the low presence of pollinators at the beginning of the season limiting the rate of visitation per flower or competition for other resources (nutrient, light) on a short suitable period.

## IV Eco-evolutionary dynamics under global changes

As a second step, we no longer consider the environmental window as fixed, but allow it to change to reflect possible consequences of global changes. Specifically, we allow the system to evolve toward quasi-equilibrium in a fixed environmental window (blue in fig 4a). We then modify environmental conditions and study the consequences of this modification for eco-evolutionary dynamics and for the maintenance of diversity. Modifications correspond to changes in the onset and ending of the environmental window (parameter j_1/2_) (eg, milder conditions in early spring and late autumn) and to a depletion of up to 90% within summer (second part of equation 4) to reflect harsher conditions mid-summer (drought, extreme temperatures, (Spinoni et al. 2018)) (fig 4a). This change consists in a linear evolution of the parameters of the environmental window for the time period *T_ch_*. We keep the integral under the curve constant, so that total environmental quality *per se* does not change in these simulations.

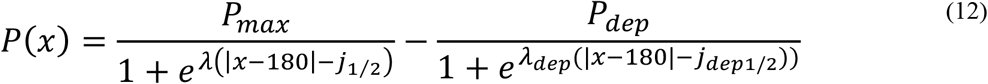

**Figure 4 :**
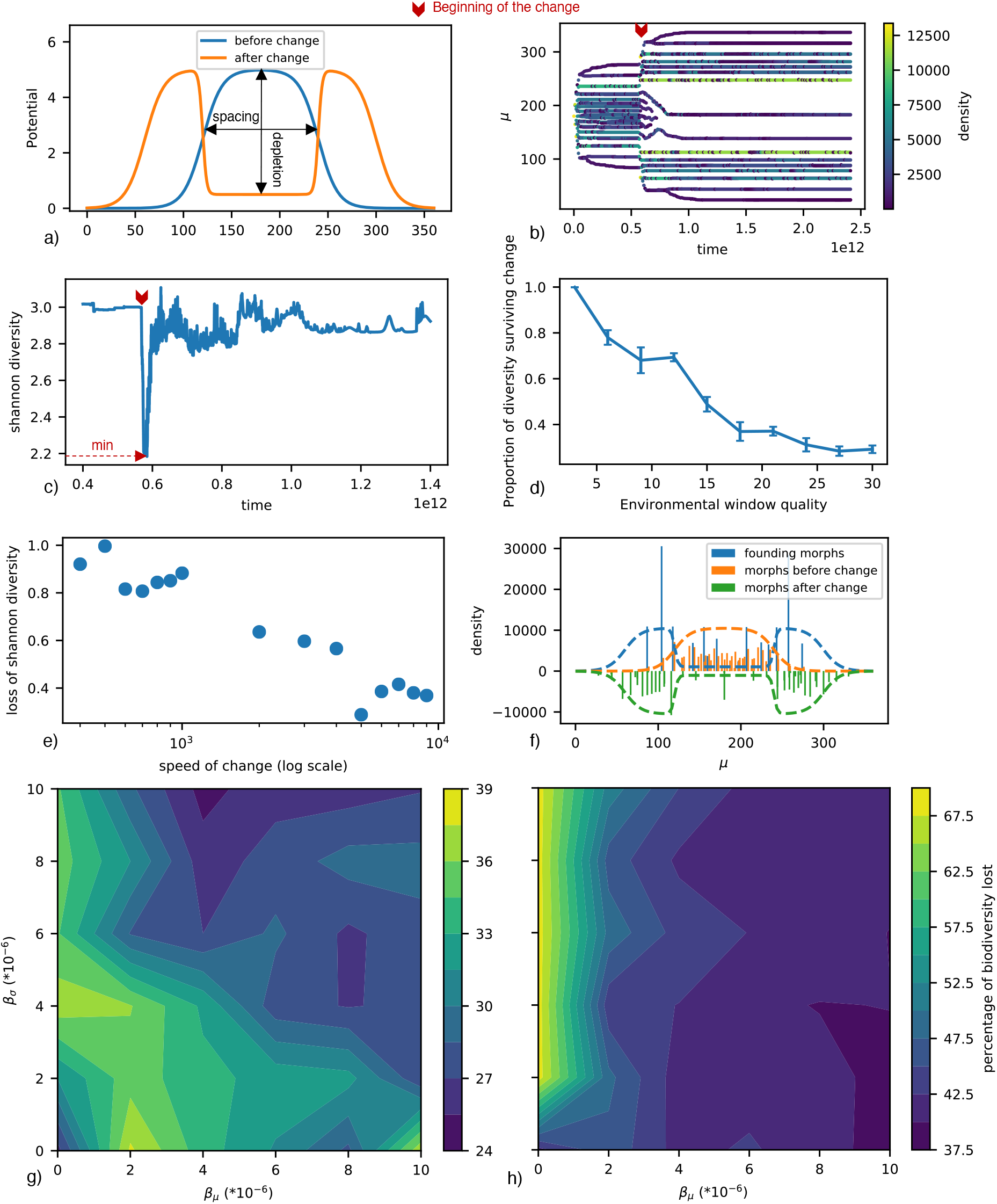
Consequences of environmental for community eco-evolutionary dynamics a) Modeling environmental changes through modifications of the environmental window b) an example of coevolutionary dynamics of mean dates under environmental changes. Densities of the phenologies are reported through variations in colors. c) Diversity dynamics in time. d) Proportion of the phenologies that contribute to diversity post change depends on the quality of the environmental window. e) Percentage of diversity loss as a function of the rate of change. Loss is measured based on the difference between Shannon diversity at the onset of environmental change and the minimum of Shannon diversity following it. (*T_ch_* ∈ [40:9000]) f) The funding phenotypes: dashed lines represent the environmental windows. Bars are the densities of each phenotype with their associated mean date. The orange bars are the ones before the change, the green ones are the phenotypes after the change. In bleu are the funding phenotypes, the density associated is the cumulated density of all its descendants. g,h) biodiversity loss (in percentage) as a function of evolutionary capacity for *μ* and *σ* in a rich (g) and in a poor environment (h). (*P_max_* = 12 for panels (bcdfg), *P_max_* = 8 for panel (h), β_*μ*_, β_σ_ = 10^−5^ for panels (bcdef), *T_ch_* = 2000 for panels (bcdfgh), *j*_1/2_ = 120 for panels (ab), *j*_1/2_ = 120 for panels (abcdefgh), *j*_1/2*dep*_ = 60 for panels (abcdefgh))

### IV.1 Evolutionary rescue of phenologies

Based on evolutionary rescue (Gomulkiewicz and Holt 1995) we expect that a given phenology may more easily persist when its evolutionary potential (here mutation rate and amplitude) is larger, and when the disturbance is less intense. Evolutionary rescue theory is however based on single species models and may not hold in complex communities where indirect effects occur (Loeuille 2019). Surprisingly, we found that the tenets of evolutionary rescue still apply to our communities of competing phenologies. Particularly, biodiversity is saved when evolution is allowed to be faster and disturbances are slower.

We vary the environmental window for different time periods *T_ch_* to simulate various speeds of current changes (fig 4e). This speed of the environmental change is to be contrasted with the adaptation capacity of phenologies, that is also systematically manipulated (fig 4gh). Finally, we investigate whether consequences of global changes are similar in poor (fig 4h) vs rich environments (fig 4g).

Following the environmental change, we observe that phenologies evolve toward newly suitable early and late dates, while most go extinct mid-year due to the depletion (Fig 4b). As expected from previous results (eg, equation 9), mid-year phenologies have large spreads (light blue, sup fig 1) as they evolve in a deteriorated environment (summer depletion).

In spite of this continuous adaptation of phenologies, modeled changes have large impacts on the maintenance of diversity. Diversity decreases drastically during the environmental change, then partially recovers (Fig4c). Diversity loss is quantified based on the difference between the diversity level observed just before the change and the minimum diversity observed post-change. We confirm the intuition that the faster the change, the greater diversity losses (Fig4e).

Fast evolution of the two traits allows a better resistance of the community (fig 4g,h). In poor environments (Fig4h), fast evolution of mean dates *μ* is particularly important. Evolution of spread *σ* alone is detrimental as phenologies adapt to the local environmental quality thereby widening and increasing competition with surrounding phenologies (low β_*μ*_ on Fig 4g,h). Mean date evolution allows phenologies to move to new suitable parts of the environmental window. In rich environments, however, evolutionary potentials of both traits jointly favor the maintenance of diversity (Fig 4g), emphasizing a stronger interaction between the two dimensions of phenologies. A possibility is that in such rich environments selection of larger spread (hence increased competition) is less prevalent.

Phenologies existing pre-change contribute differently to biodiversity post-change. Founding phenologies are mostly at the edge of the environmental window, pre-change (Fig 4f). Early and late season phenology could better maintain themselves in the face of change and drive future diversity. The proportion of initial lineages founding post-change diversity is smaller for high quality environments that have high pre-change diversity (Fig 4d). Competition has a greater effect on the loss of diversity in such systems, as they initially have many high density phenologies mid-summer. Coevolved changes in the timing and spread of phenologies in response to change then increase competition between phenologies and cause extinctions.

### IV.2 Phenology coevolution in heterogeneous space

As a final step, we extend our eco-evolutionary model to account for spatial aspects (metacommunity). This will allow us to discuss the relative contribution of spatial dynamics (dispersal) vs local evolution in determining diversity maintenance. Our model has 5 patches along a gradient of the environmental windows (latitudinal or elevation gradient, figure 5c). Patches at the lower end of the gradient have a large environmental window (eg, temperature allows activity for a large part of the year), while at the other extreme of the gradient, the environmental windows are narrower. Local ecological and evolutionary dynamics are similar to previous sections. Phenologies have a certain probability *P_m_* of dispersing to the adjacent patch every fixed time step *dt_m_*. Phenologies at the edge can only migrate inward. Once the metacommunity has settled on an eco-evolutionary dynamics quasi-equilibrium, we alter the environmental windows to mimic global changes as we did in the previous section. Before the environmental change, total diversity decreases as the dispersal rate increases (figure 5a). Such increases in dispersal homogenise the competitive context at the metacommunity scale. Consistent with previous metacommunity works, phenotypes that are better adapted to this global context are favored, while others go extinct, which explains the drop in diversity (de Mazancourt, Johnson, and Barraclough 2008; Mouquet and Loreau 2003; Thompson and Fronhofer 2019).

**Figure 5 :**
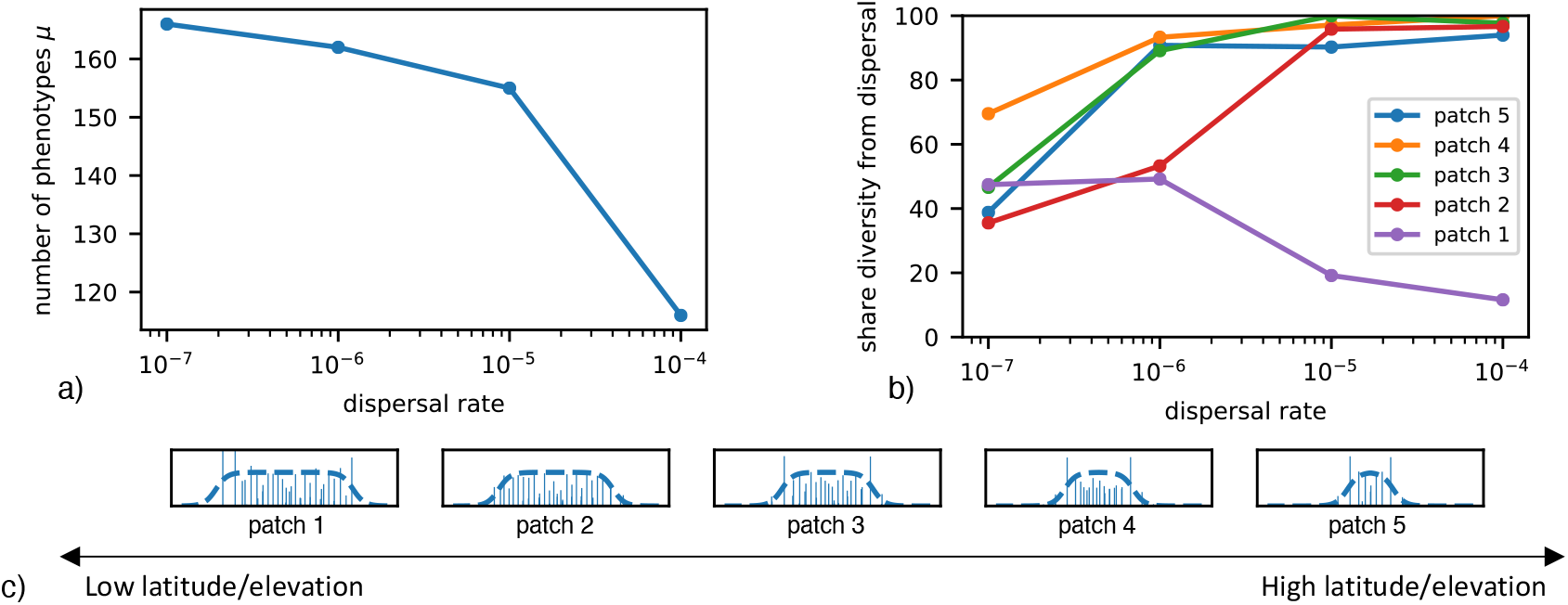
Implications of dispersal for metacommunity eco-evolutionary dynamics a) Global diversity at the evolutionary equilibrium decreases with dispersal rate. b) Share of final (post-change) diversity attributable to dispersal versus local evolution for the 5 patches c) Gradient of environmental windows (thick dashed lines) and evolved phenologies (light blue) from low latitudes (left) to high latitudes (right). (*j*_1/2_ ∈ [60,80,100,120,140] for (ab), *j*_1/2*dep*_ ∈ [30,40,60,70] for (b))

We now turn to global change simulations and study whether phenologies maintained after change come from another patch (dispersal contribution) vs *in situ* evolution (adaptation contribution). At higher dispersal rates, the share of phenotypes resulting from dispersal increases in high latitude/elevation patches 2,3,4 or 5 but decreases in the low latitude/elevation patch 1 (Figure 5b). Phenologies that could have evolved on patches 2,3,4 to fill the new environmental window have been outcompeted by phenotypes coming from lower latitudes. On patch 1 (low latitude), no pre-adapted disperser exists so that the maintenance of diversity largely rely on adaptation (Figure 5b).

## VII Discussion

Our simple model suggests that eco-evolutionary dynamics of competing phenologies often allow the emergence of diversification, leading to communities whose diversity is constrained by the available environmental window. Diversity may be partly maintained given environmental changes due to evolutionary rescue, though competitive exclusion may be intensified in some instances. The spatial model reveals that this maintenance of diversity through local evolution is dominant at low latitudes or elevations, but that dispersal contribution is dominant in other conditions.

For a fixed environment, we find that phenologies adapt to local environmental quality through changes in the phenological spread while limiting similarity leads to partitioning of the environmental window through evolution of different mean dates. Coevolution of the two traits thereby lead to a robust pattern where long phenologies are favored early and late in the year, while central phenologies are short. The empirical pattern we observe in the Swedish flora is consistent with this prediction. Among these flowering plants likely sharing roughly similar environmental windows, we find short flowering blooms during summer and longer flowering periods at the beginning and the end of the year. Such variations may be related to the abundance of pollinators, nutrient or light allowing for shorter flowering periods when environments are more suitable during summer (Kehrberger and Holzschuh 2019). Short mass blooming then increases the frequency of visits by pollinators (Ashman and Shoen 1996). On the contrary, at the beginning of the season phenologies are more spread out. Such a spread, selected in the model by competition in poor resource conditions, could be further enhanced in nature as bet hedging strategy given the risks of an uncertain pollination (Kehrberger and Holzschuh 2019; Rathcke and Lacey 1985). We want to stress that the study of flowering phenology in Sweden is here provided as an illustration, a way to test one of the key results of the model. We do not claim that such patterns will inevitably be found in nature. Among pollinators in Corsica (sup fig 2), phenological spread increases linearly with the average phenology date (Menegus 2018). One possibility is that the shape of the environmental window differs in these locations. For instance, vegetation environmental windows seem to vastly differ across Europe (Stöckli and Vidale 2004). In this article, authors observed very abundant vegetation at the beginning of the season. Insect pollinators can be expected to have a reduced phenology early in the year when vegetation is flourishing. However, in our model, the minimum limit of phenological spread only depends on the environmental richness, whereas this limit certainly also depends on physiological constraints related to development. The phenologies can then be artificially infinitely fine if the environment is infinitely rich. Some models explicitly take into account organism development as a function of the environment (Johansson and Bolmgren 2019). This sets a more realistic minimum limit for the phenological spread, but would not necessarily change our predictions.

Next to the hypothesis of variation in the shape of the environmental window, we want to stress that the pattern we propose here is a signature of the coevolution of competing phenologies. We do not expect to observe it when phenologies are mainly constrained not by interspecific interactions, but directly by abiotic conditions. For instance, if late phenologies were to be mostly determined by abiotic factors, competition playing a lesser role (few active individuals), while competition decrease progressively in the year, we would expect an increased phenological spread only late in the year, which would be consistent with the pattern here reported in Corsica.

Our model often allows diversification, following a partitioning of the environmental window that leads to a limiting similarity of phenologies, thereby reducing competition (Macarthur and Levins 1967). Such a temporal partitioning of activity of species competing for common resources has been reported in various contexts. Mediterranean amphibians share the use of ponds for breeding (Richter-Boix, Llorente, and Montori 2006). Our competition term would then correspond to competition for spawning space. In flowering plants, many examples of exclusion through competition for pollinators have been shown (Willmer 2011). For systems in which hybrids have low fitness (Waser 1978), competition for pollinator exclusivity may cause the partitioning of phenologies. Conversely, having very distinct phenologies could be indicative of limiting competition. The diversification process we report here maintains various levels of diversity. We find that diversity mainly increases with the width of the environmental window when spread is fixed and only phenological dates evolve. In case of coevolution, diversity is not tightly linked to the width of the environmental window, but rather increases the total quality of the environment. The difference has be explained by the fact that high quality environments select for narrower spread, so that if spread can evolve, more phenologies can be packed within the environmental window.

While current changes threaten biodiversity, evolution may play a key role in saving certain species (evolutionary rescue, (Gomulkiewicz and Holt 1995)). It is however unclear whether such a positive role of evolution may remain in a community context (Johansson 2008; Loeuille 2019; Osmond and de Mazancourt 2013). For instance, in a competition context, fast evolution of a phenology may allow it to outcompete neighbor phenologies. Theoretically, for a given phenology, evolutionary rescue may happen due to changes on either dimensions: changing dates to follow suitable conditions, or changing spread to cover more conditions. Interestingly, we find that adaptive evolution tracks the new shape of the environmental window mainly through the evolution of the mean date, particularly in poor environments. Shifts in reproductive phenologies of plants in California towards earlier and later dates in the year are similar to patterns observed in our model (Parmesan 2006; Sherry et al. 2007). Phenological shifts of dates have been linked to adaptive evolution for plants (Franks, Sim, and Weis 2007), for amphibians (Phillimore et al. 2010) and for pollinators (Duchenne et al. 2020). If this evolution is fast enough to match the speed of change, we observe an evolutionary rescue of part of our community. Evolution of phenological spread can in some instances lead to increased competition and diversity losses. This result is consistent with earlier results emphasizing that competition can alter evolutionary rescue. (Johansson 2008). Note however that competition possibly facilitates evolutionary rescue when it sufficiently increases selection in the direction of environmental changes (Osmond and de Mazancourt 2013).

Increasing competition through the evolution of the date or spread may decrease diversity. Increases of overlap of phenologies can come from antagonistic forces selecting them. Asymmetries in the evolutionary potential of the different species of the community may increase phenology overleap as well, potentially leading to competitive exclusion (THACKERAY et al. 2010; Visser, te Marvelde, and Lof 2012). Evolutionary potentials may be related to variations in population size, generation time or mutation rates (Frankham 1996). Changes in phenology overlap has been reported in Texas amphibian communities possibly leading to increased competition (Carter, Saenz, and Rudolf 2018). In a changing environment, when species adapt while competing, some often end up outcompeting the other, partly by preventing its evolution as competition lowers population sizes and corresponding evolutionary potentials (Johansson 2008; Norberg et al. 2012). While these phenomena are well known in a spatial context (Norberg et al. 2012; Thompson and Fronhofer 2019), we propose that they also likely happen for phenological shifts.

Whether rescue of biodiversity through niche tracking may happen through dispersal or adaptation depends on geographical position as well as the ability to disperse. Our model suggests that at high connectivity, dispersal plays a dominant role for niche tracking compared to evolutionary contribution. Increasing dispersal allows a fast redistribution of pre-adapted phenologies more quickly, consistent with observations of a more general model (Thompson and Fronhofer 2019). On the other hand, if no pre-adapted phenotype exists, dispersal has a negative contribution. In our simulations, this happens for the low altitude/elevation patch, where the role of evolution is dominant. The dominant role of evolution at low latitudes is consistent with another (non phenology) model (Norberg et al. 2012). Consistent with this view, empirical analyses of the role of evolution in extremely hot environments have been undertaken. For instance, (Sinervo et al. 2010) particularly studied Mexican lizard communities and proposed that by 2050 many areas could lose all their lizard diversity and favor only extremophiles (Sinervo et al. 2010). This work also suggests that the evolutionary potential of these populations may be too limited to allow evolutionary rescue. Because in other latitudes a dominant role of dispersal is found, this raises the question of connectivity. Habitat fragmentation could affect the ability of species to disperse, creating genetic isolation and decreasing colonization potentials (Clark, Lewis, and Horvath 2001; Jump and Penuelas 2005). Some places like islands or the high mountain floors cannot rely on dispersal and can only count on adaptive evolution (Krajick 2004; Thuiller et al. 2005). The ability of species to disperse is therefore likely to vary with habitat.

The model we propose here is very simple and therefore has many limitations. Particularly, the diversification process is linked to species competition, so that it is necessary that competition plays an important structuring role for the model to apply. While competition is key here, many other mechanisms could of course allow the emergence and maintenance of diversity, from the specialization on a precise set of conditions that would limit the temporal presence of a species, to the evolution of bet-hedging strategies in a (within year) varying environment. Our model is however an invitation to consider the possible role of interspecific interactions (here competition) to better understand the effect of current changes on the eco-evolutionary dynamics of natural communities.

## Acknowledgements

We thank Avril Weinbach and Ove Ericsson for data collection in the herbarium of Stockholm. We thank We also thank the ANR for funding this research. We thanks Giulio Menegus and Adrien Perrard for their data collection of pollinators in Corsica and their discussion on phenological patterns.

---

**Supplementary figure 1 :**
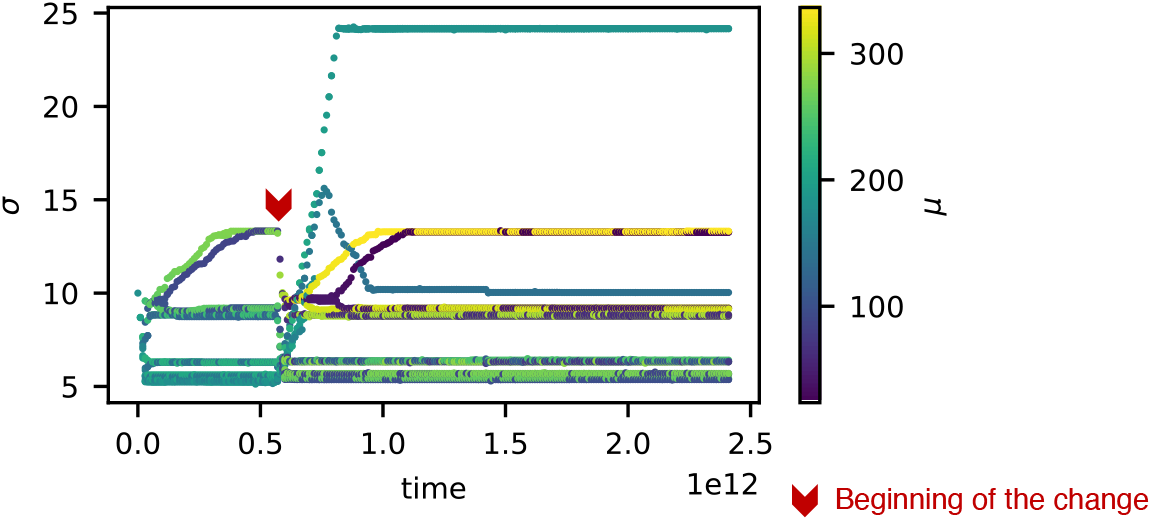
An example of coevolutionary dynamics of spread under environmental changes. Variation of colors shows the associated mean date.

**Supplementary figure 2 :**
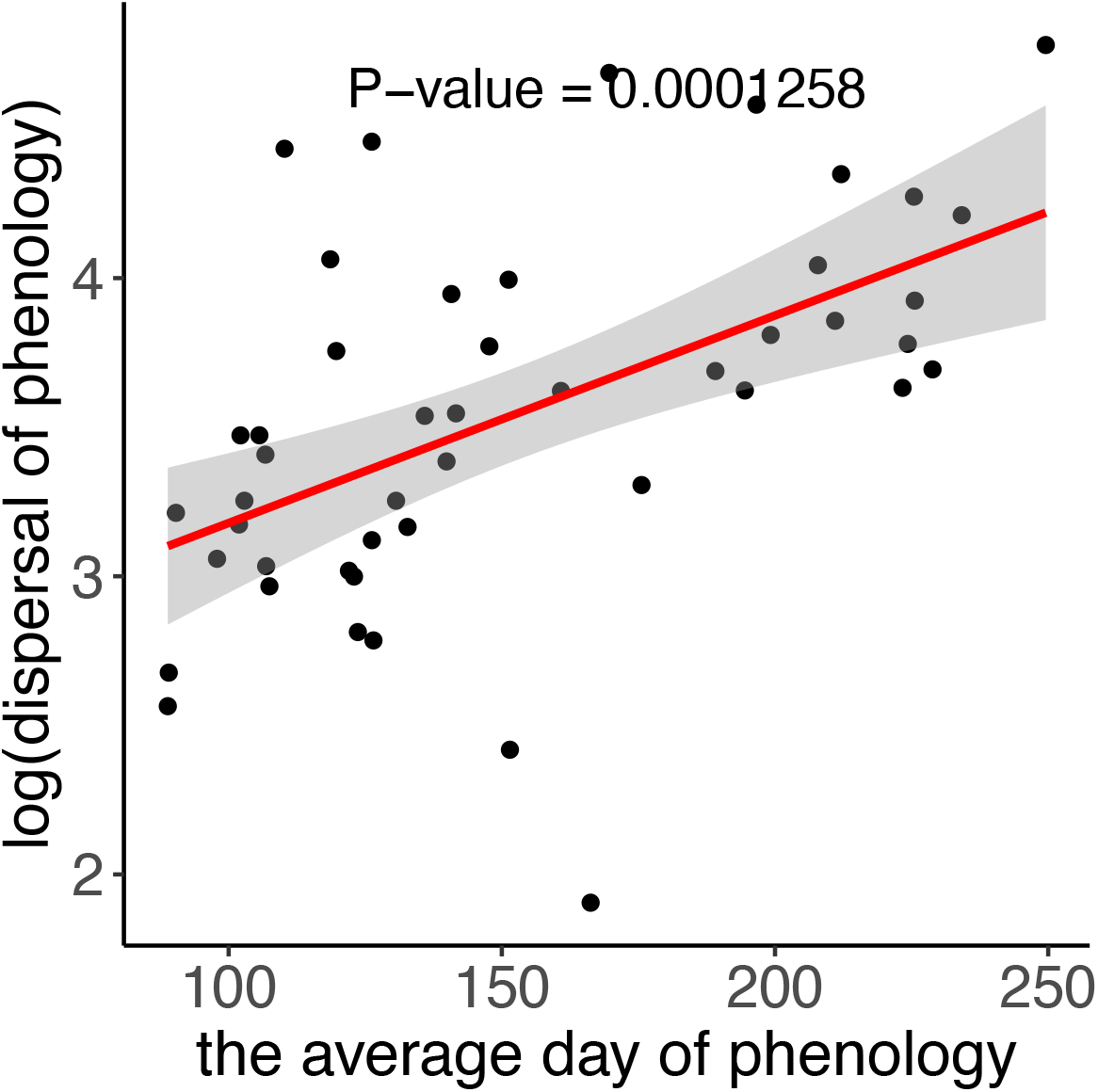
Distribution of the mean date of phenology and their spread along the year for pollinators of Corsica (Menegus 2018). The best fit is a linear regression model. The methods used are the same that the one of figure 3c.

## Computation of the singularity for *σ*

It is assumed that losses are minimal when the maximum of the Gaussian fits the environmental window.

Let’s take equation of the exploitation of the environment over time (2) :

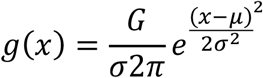

The maximum of this function if for 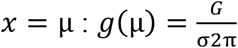

At this date the richness of the environment is *P*(*μ*), under the assumption of relatively low spread and not too close from the edge of the window:

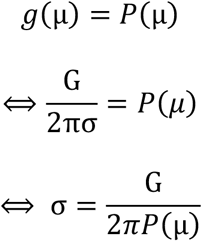

## Notes

### Competing Interest Statement

The authors have declared no competing interest.

## References

Alford, Ross A, and Henry M Wilbur. 1985. “Priority Effects in Experimental Pond Communities: Competition between Bufo and Rana.” Ecology 66 (4): 1097–1105. https://doi.org/10.2307/1939161.

Appanah, S. 1985. “General Flowering in the Climax Rain Forests of South-East Asia.” Journal of Tropical Ecology 1 (3): 225–40. https://doi.org/10.1017/S0266467400000304.

Ashman, T. L., and D. J. Shoen. 1996. Floral Biology. Edited by David G. Lloyd and Spencer C. H. Barrett. CHAPMAN &. Boston, MA: Springer US. https://doi.org/10.1007/978-1-4613-1165-2.

Augspurger, Carol K. 1981. “Reproductive Synchrony of a Tropical Shrub: Experimental Studies on Effects of Pollinators and Seed Predators in Hybanthus Prunifolius (Violaceae).” Ecology 62 (3): 775–88. https://doi.org/10.2307/1937745.

Botes, Christo, Steven D. Johnson, and Richard M. Cowling. 2008. “Coexistence of Succulent Tree Aloes: Partitioning of Bird Pollinators by Floral Traits and Flowering Phenology.” Oikos 117 (6): 875–82. https://doi.org/10.1111/j.0030-1299.2008.16391.x.

Carter, Shannon K., Daniel Saenz, and Volker H. W. Rudolf. 2018. “Shifts in Phenological Distributions Reshape Interaction Potential in Natural Communities.” Edited by Andrew Sih. Ecology Letters 21 (8): 1143–51. https://doi.org/10.1111/ele.13081.

Chesson, Peter. 2000. “Mechanisms of Maintenance of Species Diversity.” Annual Review of Ecology and Systematics 31 (1): 343–66. https://doi.org/10.1146/annurev.ecolsys.31.1.343.

Chuine, Isabelle, and Jacques Régnière. 2017. “Process-Based Models of Phenology for Plants and Animals.” Annual Review of Ecology, Evolution, and Systematics 48 (1): 159–82. https://doi.org/10.1146/annurev-ecolsys-110316-022706.

Clark, James S, Mark Lewis, and Lajos Horvath. 2001. “Invasion by Extremes: Population Spread with Variation in Dispersal and Reproduction.” The American Naturalist 157 (5): 537–54. https://doi.org/10.1086/319934.

Dieckmann, Ulf, and Richard Law. 1996. “The Dynamical Theory of Coevolution: A Derivation from Stochastic Ecological Processes.” Journal of Mathematical Biology 34 (5–6): 579–612. https://doi.org/10.1007/BF02409751.

Duchenne, F, E Thébault, D Michez, M Elias, M Drake, M Persson, J. S. Rousseau-Piot, M Pollet, P Vanormelingen, and C Fontaine. 2020. “Phenological Shifts Alter the Seasonal Structure of Pollinator Assemblages in Europe.” Nature Ecology & Evolution 4 (1): 115–21. https://doi.org/10.1038/s41559-019-1062-4.

Eshel, Ilan. 1983. “Evolutionary and Continuous Stability.” Journal of Theoretical Biology 103 (1): 99–111. https://doi.org/10.1016/0022-5193(83)90201-1.

Frankham, Richard. 1996. “Relationship of Genetic Variation to Population Size in Wildlife.” Conservation Biology 10 (6): 1500–1508. https://doi.org/10.1046/j.1523-1739.1996.10061500.x.

Franks, S. J., S. Sim, and A. E. Weis. 2007. “Rapid Evolution of Flowering Time by an Annual Plant in Response to a Climate Fluctuation.” Proceedings of the National Academy of Sciences 104 (4): 1278–82. https://doi.org/10.1073/pnas.0608379104.

Geritz, S.A.H., É. Kisdi, G. Meszéna, and J.A.J. Metz. 1998. “Evolutionarily Singular Strategies and the Adaptive Growth and Branching of the Evolutionary Tree.” Evolutionary Ecology 12 (1): 35–57. https://doi.org/10.1023/A:1006554906681.

Gomulkiewicz, Richard, and Robert D Holt. 1995. “When Does Evolution by Natural Selection Prevent Extinction?” Evolution 49 (1): 201. https://doi.org/10.2307/2410305.

Guisan, Antoine, and Wilfried Thuiller. 2005. “Predicting Species Distribution: Offering More than Simple Habitat Models.” Ecology Letters 8 (9): 993–1009. https://doi.org/10.1111/j.1461-0248.2005.00792.x.

Husby, Arild, Marcel E. Visser, and Loeske E. B. Kruuk. 2011. “Speeding Up Microevolution: The Effects of Increasing Temperature on Selection and Genetic Variance in a Wild Bird Population.” Edited by Joel G. Kingsolver. PLoS Biology 9 (2): e1000585. https://doi.org/10.1371/journal.pbio.1000585.

Johansson, Jacob. 2008. “EVOLUTIONARY RESPONSES TO ENVIRONMENTAL CHANGES: HOW DOES COMPETITION AFFECT ADAPTATION?” Evolution 62 (2): 421–35. https://doi.org/10.1111/j.1558-5646.2007.00301.x.

Johansson, Jacob, and Kjell Bolmgren. 2019. “Is Timing of Reproduction According to Temperature Sums an Optimal Strategy?” Ecology and Evolution 9 (20): 11598–605. https://doi.org/10.1002/ece3.5601.

Jonzen, N. 2006. “Rapid Advance of Spring Arrival Dates in Long-Distance Migratory Birds.” Science 312 (5782): 1959–61. https://doi.org/10.1126/science.1126119.

Jump, Alistair S., and Josep Penuelas. 2005. “Running to Stand Still: Adaptation and the Response of Plants to Rapid Climate Change.” Ecology Letters 8 (9): 1010–20. https://doi.org/10.1111/j.1461-0248.2005.00796.x.

Kehrberger, Sandra, and Andrea Holzschuh. 2019. “How Does Timing of Flowering Affect Competition for Pollinators, Flower Visitation and Seed Set in an Early Spring Grassland Plant?” Scientific Reports 9 (1): 15593. https://doi.org/10.1038/s41598-019-51916-0.

Krajick, Kevin. 2004. “CLIMATE CHANGE: All Downhill From Here?” Science 303 (5664): 1600–1602. https://doi.org/10.1126/science.303.5664.1600.

Kubisch, Alexander, Tobias Degen, Thomas Hovestadt, and Hans Joachim Poethke. 2013. “Predicting Range Shifts under Global Change: The Balance between Local Adaptation and Dispersal.” Ecography 36 (8): 873–82. https://doi.org/10.1111/j.1600-0587.2012.00062.x.

Lloyd, J, and J A Taylor. 1994. “On the Temperature Dependence of Soil Respiration.” Functional Ecology 8 (3): 315. https://doi.org/10.2307/2389824.

Loeuille, Nicolas. 2019. “Eco-Evolutionary Dynamics in a Disturbed World: Implications for the Maintenance of Ecological Networks.” F1000Research 8 (0): 97. https://doi.org/10.12688/f1000research.15629.1.

MacArthur, R., and R. Levins. 1964. “COMPETITION, HABITAT SELECTION, AND CHARACTER DISPLACEMENT IN A PATCHY ENVIRONMENT.” Proceedings of the National Academy of Sciences 51 (6): 1207–10. https://doi.org/10.1073/pnas.51.6.1207.

Macarthur, Robert, and Richard Levins. 1967. “The Limiting Similarity, Convergence, and Divergence of Coexisting Species.” The American Naturalist 101 (921): 377–85. https://doi.org/10.1086/282505.

Mazancourt, C. de, E. Johnson, and T. G. Barraclough. 2008. “Biodiversity Inhibits Species’ Evolutionary Responses to Changing Environments.” Ecology Letters 11 (4): 380–88. https://doi.org/10.1111/j.1461-0248.2008.01152.x.

Menegus, Giulio. 2018. “Seasonal Morphological Variation in a Pollinator Community: The Wild Bees of Southern Corsica.” UNIVERSITÀ DEGLI STUDI DI PADOVA.

Metz, J.A.J., R.M. Nisbet, and S.A.H. Geritz. 1992. “How Should We Define ‘Fitness’ for General Ecological Scenarios?” Trends in Ecology & Evolution 7 (6): 198–202. https://doi.org/10.1016/0169-5347(92)90073-K.

Mitchell, Randall J, Rebecca J Flanagan, Beverly J Brown, Nickolas M Waser, and Jeffrey D Karron. 2009. “New Frontiers in Competition for Pollination.” Annals of Botany 103 (9): 1403–13. https://doi.org/10.1093/aob/mcp062.

Mouquet, Nicolas, and Michel Loreau. 2003. “Community Patterns in Source-Sink Metacommunities.” The American Naturalist 162 (5): 544–57. https://doi.org/10.1086/378857.

Norberg, Jon, Mark C. Urban, Mark Vellend, Christopher A. Klausmeier, and Nicolas Loeuille. 2012. “Eco-Evolutionary Responses of Biodiversity to Climate Change.” Nature Climate Change 2 (10): 747–51. https://doi.org/10.1038/nclimate1588.

Nussey, Daniel H. 2005. “Selection on Heritable Phenotypic Plasticity in a Wild Bird Population.” Science 310 (5746): 304–6. https://doi.org/10.1126/science.1117004.

Osmond, Matthew Miles, and Claire de Mazancourt. 2013. “How Competition Affects Evolutionary Rescue.” Philosophical Transactions of the Royal Society B: Biological Sciences 368 (1610): 20120085. https://doi.org/10.1098/rstb.2012.0085.

Parmesan, Camille. 2006. “Ecological and Evolutionary Responses to Recent Climate Change.” Annual Review of Ecology, Evolution, and Systematics 37 (1): 637–69. https://doi.org/10.1146/annurev.ecolsys.37.091305.110100.

Parmesan, Camille, and Gary Yohe. 2003. “A Globally Coherent Fingerprint of Climate Change Impacts across Natural Systems.” Nature 421 (6918): 37–42. https://doi.org/10.1038/nature01286.

Phillimore, A. B., J. D. Hadfield, O. R. Jones, and R. J. Smithers. 2010. “Differences in Spawning Date between Populations of Common Frog Reveal Local Adaptation.” Proceedings of the National Academy of Sciences 107 (18): 8292–97. https://doi.org/10.1073/pnas.0913792107.

Primack, Richard, and Elizabeth Stacy. 1998. “Cost of Reproduction in the Pink Lady’s Slipper Orchid (Cypripedium Acaule, Orchidaceae): An Eleven-Year Experimental Study of Three Populations.” American Journal of Botany 85 (12): 1672–79. https://doi.org/10.2307/2446500.

Rathcke, Beverly, and Elizabeth P Lacey. 1985. “Phenological Patterns of Terrestrial Plants.” Annual Review of Ecology and Systematics 16 (1): 179–214. https://doi.org/10.1146/annurev.es.16.110185.001143.

Richter-Boix, Alex, Gustavo A. Llorente, and Albert Montori. 2006. “Breeding Phenology of an Amphibian Community in a Mediterranean Area.” Amphibia-Reptilia 27 (4): 549–59. https://doi.org/10.1163/156853806778877149.

Sherry, Rebecca A, Xuhui Zhou, Shiliang Gu, J. A. Arnone, David S Schimel, Paul S Verburg, Linda L Wallace, and Yiqi Luo. 2007. “Divergence of Reproductive Phenology under Climate Warming.” Proceedings of the National Academy of Sciences 104 (1): 198–202. https://doi.org/10.1073/pnas.0605642104.

Sinervo, Barry, F. Mendez-de-la-Cruz, Donald B. Miles, Benoit Heulin, Elizabeth Bastiaans, M. Villagran-Santa Cruz, Rafael Lara-Resendiz, et al. 2010. “Erosion of Lizard Diversity by Climate Change and Altered Thermal Niches.” Science 328 (5980): 894–99. https://doi.org/10.1126/science.1184695.

Snow, D. W. 1965. “A Possible Selective Factor in the Evolution of Fruiting Seasons in Tropical Forest.” Oikos 15 (2): 274. https://doi.org/10.2307/3565124.

Spinoni, Jonathan, Jürgen V Vogt, Gustavo Naumann, Paulo Barbosa, and Alessandro Dosio. 2018. “Will Drought Events Become More Frequent and Severe in Europe?” International Journal of Climatology 38 (4): 1718–36. https://doi.org/10.1002/joc.5291.

Stöckli, R., and P. L. Vidale. 2004. “European Plant Phenology and Climate as Seen in a 20-Year AVHRR Land-Surface Parameter Dataset.” International Journal of Remote Sensing 25 (17): 3303–30. https://doi.org/10.1080/01431160310001618149.

Thackeray, STEPHEN J., TIMOTHY H. Sparks, MORTEN Frederiksen, SARAH Burthe, PHILIP J. Bacon, JAMES R. Bell, MARC S. Botham, et al. 2010. “Trophic Level Asynchrony in Rates of Phenological Change for Marine, Freshwater and Terrestrial Environments.” Global Change Biology 16 (12): 3304–13. https://doi.org/10.1111/j.1365-2486.2010.02165.x.

Thompson, Patrick L, and Emanuel A Fronhofer. 2019. “The Conflict between Adaptation and Dispersal for Maintaining Biodiversity in Changing Environments.” Proceedings of the National Academy of Sciences 116 (42): 21061–67. https://doi.org/10.1073/pnas.1911796116.

Thuiller, Wilfried, Sandra Lavorel, M. B. Araujo, M. T. Sykes, and I. C. Prentice. 2005. “Climate Change Threats to Plant Diversity in Europe.” Proceedings of the National Academy of Sciences 102 (23): 8245–50. https://doi.org/10.1073/pnas.0409902102.

Tylianakis, Jason M., Raphael K. Didham, Jordi Bascompte, and David A. Wardle. 2008. “Global Change and Species Interactions in Terrestrial Ecosystems.” Ecology Letters 11 (12): 1351–63. https://doi.org/10.1111/j.1461-0248.2008.01250.x.

Visser, Marcel E, Luc te Marvelde, and Marjolein E. Lof. 2012. “Adaptive Phenological Mismatches of Birds and Their Food in a Warming World.” Journal of Ornithology 153 (S1): 75–84. https://doi.org/10.1007/s10336-011-0770-6.

Vitousek, Peter M. 1992. “Global Environmental Change: An Introduction.” Annual Review of Ecology and Systematics 23 (1): 1–14. https://doi.org/10.1146/annurev.es.23.110192.000245.

Waser, Nickolas M. 1978. “Interspecific Pollen Transfer and Competition between Co-Occurring Plant Species.” Oecologia 36 (2): 223–36. https://doi.org/10.1007/BF00349811.

Waser, Nickolas M. 1979. “Pollinator Availability as a Determinant of Flowering Time in Ocotillo (Fouquieria Splendens).” Oecologia 39 (1): 107–21. https://doi.org/10.1007/BF00346001.

Weinbach, Avril. 2015. “Relationships between Flowering Time and Distribution Ranges of Northern European Plants.”

Wheelwright, Nathaniel T. 1985. “Competition for Dispersers, and the Timing of Flowering and Fruiting in a Guild of Tropical Trees.” Oikos 44 (3): 465. https://doi.org/10.2307/3565788.

Willmer, Pat. 2011. Pollination and Floral Ecology. Princeton University Press. https://doi.org/10.1515/9781400838943.

